# Unconstrained dosing agar (UDA) Reduces Stress in Mouse Oral Administration

**DOI:** 10.64898/2026.02.25.707935

**Authors:** Melissa Lee, Cornel Fraefel, Catherine Eichwald, Claudio Aguilar

## Abstract

**Background:** Oral gavage is the standard method for delivering drugs and other substances orally in rodent studies, but it can cause significant stress and risk injury. To improve animal welfare and reduce confounding stress effects, this study aimed to replace oral gavage by developing and testing a new voluntary ingestion method that is easy to adopt, minimizes stress in mice, and is suitable for a wide range of compounds.

**Results:** We developed a soft agar formulation with an appealing scent and taste that mice readily consumed without fasting or restraint. We called this method “unconstrained dosing agar” (UDA). Analysis of fecal corticosterone levels demonstrated that the method is associated with low stress in the animals. After training, mice quickly consumed the agar units. Body weight gain was unaffected by the treatment.

**Conclusions:** This study introduces a simple, low-stress method for administering substances orally in mice. By encouraging voluntary consumption and removing the need for fasting or restraint, this method provides a practical alternative to oral gavage and could improve animal welfare and experimental consistency.

## BACKGROUND

In rodent studies, physiological parameters such as heart rate, blood pressure, or corticosterone levels can be significantly elevated by external stressors, including routine handling, thereby influencing or confounding experimental measurements (1). A relevant source of stress for rodents during experiments is oral gavage, which is currently the standard method for administering drugs or xenobiotics (2). While oral gavage allows for consistent and accurate dosing in pharmaceutical or toxicological studies, it also has several drawbacks that may compromise animal welfare (3). During oral gavage, animals experience physical stress, including handling and restraint, the insertion of a rigid metal or plastic needle from the mouth directly into the stomach, potential breathing interference, and stomach distension (2). In addition to physical stress, oral gavage also poses serious risks such as accidental tracheal administration, reflux, aspiration pneumonia, esophageal impaction, trauma or perforation, hemothorax, and death (4). To address these issues in mice, alternative methods of oral substance administration in varying forms have recently been developed, including flavored dough (4), hazelnut spread (5), peanut butter (6), oat flakes (7), jam (8), different-sized jellies (9, 10) or liquid solutions (11, 12). However, these methods still have varying levels of restraint or water and food deprivation, and mice are typically trained over 3 to 5 days to voluntarily consume the substance under investigation.

Here, we aimed to develop a new formulation that combines voluntary consumption in mice with the convenience of easy administration and flexible dosing, thereby promoting animal welfare. We refer to this formulation and method as <u>u</u>nconstrained <u>d</u>ose <u>a</u>dministration (UDA). The formulation is based on a soft agar unit containing condensed milk and bacon extract, which the mice voluntarily eat without fasting, restraint, or manipulation. Since UDA is based on a semi-solid matrix, it works well with compounds that have low water solubility without needing detergents. The levels of stress caused by UDA were measured through fecal corticosterone concentrations and compared with those from MDA, a recently described low-stress oral administration method (11). Our results suggest that UDA is easy to implement, induces low levels of stress, does not affect weight gain, and, as such, represents a viable alternative to oral gavage in laboratory mice.

## METHODS

### Ethics statement

All mouse experiments were performed in accordance with the guidelines of the Swiss federal government’s animal experimentation law (SR 455.163; TVV). The Cantonal Veterinary Office of Zurich, Switzerland, approved the protocols under animal experimentation number ZH056/2024.

### Animals

Male and female C57BL/6 were obtained from Charles River Laboratories (Sulzfeld, Germany) at the age of 6 weeks. Mice were housed in individually ventilated cages (IVC type 2 long) in groups of 3 animals of the same sex and supplied with standard enrichment (bedding, transparent red house, tissues and crinklets). Animals within a cage were assigned the same experimental condition (MDA, UDA or control). Throughout the experiment, the cages were kept at controlled temperature (22.5 ± 1.5°C) and humidity (50 ± 10%) with a 14/10 light cycle (lights off: 8:00 PM to 6:00 AM). Animals had *ad libitum* access to standard rodent chow (Kliba Nafag diet 3335, Kaiseraugst, Switzerland) and water. For all experimental conditions, the animals were handled by two equally trained and experienced researchers. Animals were handled exclusively using the house as support and were never lifted by the tail. Upon ending the experiment, the animals were not euthanized but instead were placed in a rehoming program from the University of Zürich. (https://www.uzh.ch/en/researchinnovation/ethics/animals/3R-replace-reduce-refine/rehoming.html).

### Unconstrained dose administration (UDA) procedure

The animals were presented with 400 µl agar-based units (see below) and were trained in groups (without isolation) during two consecutive days with one training session per day to overcome the innate neophobia of the mice towards novel food sources (13). Access to food or water was not interrupted before or during the training sessions. On the first and second training days, one agar-based unit per animal was introduced into the cage, using the house inverted as support. The animals were video-recorded until complete consumption, and the elapsed time was. registered. Immediately after the animals consumed the units, the house was inverted again to its normal position. On the third day, the animals were isolated in individual cages, with one agar-based unit presented per animal in the same way as during training. The animals were regrouped after 24 hours, and feces were collected for further analysis.

### Micropipette-guided drug administration (MDA) procedure

We compared our results with those obtained with the MDA method. The MDA training and administration were performed as previously described (11, 12), using a diluted condensed milk dose of 2 ml/kg per animal over two days with one training session per day. The animals were video-recorded, and the time required for administration was registered. After the MDA procedure, the animals were isolated in individual cages for fecal collection over 24 h and then regrouped.

### Quantification of corticosterone (CORT) in feces

Fecal corticosterone metabolites (FCM) were extracted by incubating feces in 96% ethanol (WR International GmbH) at a ratio of 5 ml ethanol per gram of feces. Incubation was performed at room temperature (24± 1°C) on a rotary disc, using 15 ml sterile tubes (Sarsted AG, Germany). After overnight incubation, tubes were vortexed for 1 minute and then centrifuged at 4400 *x g* for 15 minutes at 4°C. The supernatants were collected and stored at -20°C until further use. Corticosterone levels were analysed in duplicate using a DRG-Diagnostics corticosterone ELISA kit (EIA-4164, DRG Instruments GmbH, Marburg, Germany), following the manufacturer’s instructions. The corticosterone standards included in the kit were replaced with a custom 7-point standard curve prepared with analytical corticosterone (46148, Merck) resuspended in 96% ethanol, using a concentration range from 4.5 to 288 nM. The colorimetric reaction was quantified at 450nm using a SPARK reader (Tecan, Switzerland).

### Preparation of UDA units

The UDA agar-based units were prepared using the wells of a sterile 96-well plate (F-base 92096, TPP, Switzerland). All solutions were prewarmed to 60°C before mixing. One 400 µl agar-based unit was prepared by mixing 200 µl of a 3% agar solution (A0949, ITW reagents) with a solution containing 60 µl condensed milk (Migros Kondensmilch), 5 µl ethanol-based bacon extract (see below), and 135 µl PBS pH 7.4 (Gibco, Thermo Fisher Scientific). This preparation can be scaled up to the number of UDA agar-based units required. The mix was aliquoted into the wells of the plate using a micropipette and incubated at room temperature in sterile conditions until gelification. The UDA agar-based units were removed from the 96-well plate using a sterile micro spatula (Fig. S1), collected, and stored at 4°C until further use, usually within 24 h.

### Bacon extract preparation

Bacon extract was added to the formulation to enhance the palatability and scent attraction of the agar-based units. Using a 500 ml beaker flask, 175g of bacon cubes were heated with a Bunsen burner at maximum heat for over 15 min, until the fat liquefied. The mixture was then transferred to a 1 L Erlenmeyer flask containing 200 mL of 96% ethanol and stirred at low speed for 2 h using a magnetic stirrer. To remove the solid particles, the mixture was filtered through a funnel and filter paper (Whatman No. 1) into a 250 ml centrifuge bottle (Nalgene), then incubated for 15 min at -80°C. The solution was then centrifuged at 16’000 x g for 15 min at 4°C using a GSA rotor. The supernatant was recovered, filtered through a 0.45 µm filter, and stored at 4°C for up to 2 months.

### Body weight monitoring

All animals were weighed once a day for 3 consecutive days following each experimental intervention, i.e., after training or after oral administration. Afterwards, they were monitored once a week.

### Statistical analysis

The data were analyzed using parametric analysis of variance (ANOVA), Student’s t-test, or simple linear regression as needed. Statistical significance was set at p<0.05. All analyses were performed using GraphPad Prism version 10.6.1 for macOS (GraphPad Software, Boston, Massachusetts, USA).

## RESULTS

### UDA induces low levels of stress on mice and does not impact weight gain

It has recently been shown that plasma CORT levels in mice administered orally using the MDA technique are significantly lower than those in mice subjected to oral gavage, thus revealing MDA as an interesting alternative to oral gavage for oral administration (11, 14). There are, however, a few shortcomings in the MDA technique that we have addressed in the present study, such as the need to restrain (total or light by the tail) during the training period, and that the formulation of the dose favours water-soluble compounds for administration, unless detergents are included in the formulation. We aimed to further refine the method by presenting mice with an agar unit that they will voluntarily consume without restraint, and we will hereafter refer to this methodology as <u>u</u>nconstrained <u>d</u>ose <u>a</u>dministration (UDA). Consequently, we used MDA as the reference for the low-stress method and assessed the stress levels induced by UDA. There were two training sessions over two consecutive days for both the MDA and the UDA methods. For MDA, each mouse had to be trained individually, with total restraint on the first day and lighter restraint on the second day. In the UDA case, mice were trained while grouped inside the cage on both days, with no manipulation. The untreated control group was neither trained nor manipulated. To measure and compare the stress levels induced by UDA or MDA, we opted for a non-invasive methodology, *i*.*e*., measuring fecal FCM (fecal corticosterone metabolites) levels over 24 h following each treatment (15, 16). For this purpose, mice of the UDA, MDA, and control groups were isolated in single cages, and then the treatments were applied. After 24h of isolation, the animals were regrouped, feces were collected, and fecal CORT was quantified. The treatment procedure was repeated at days 1, 21, and 42 (Fig. 1A). Our results show that fecal CORT levels were similar across the three groups, and notably, there were no significant differences in fecal CORT levels between the MDA and UDA groups throughout the treatment period (Fig. 1B-D). This suggests that the stress caused by the treatments was low and consistent across all groups, compared with the untreated control group.

**Figure 1.**
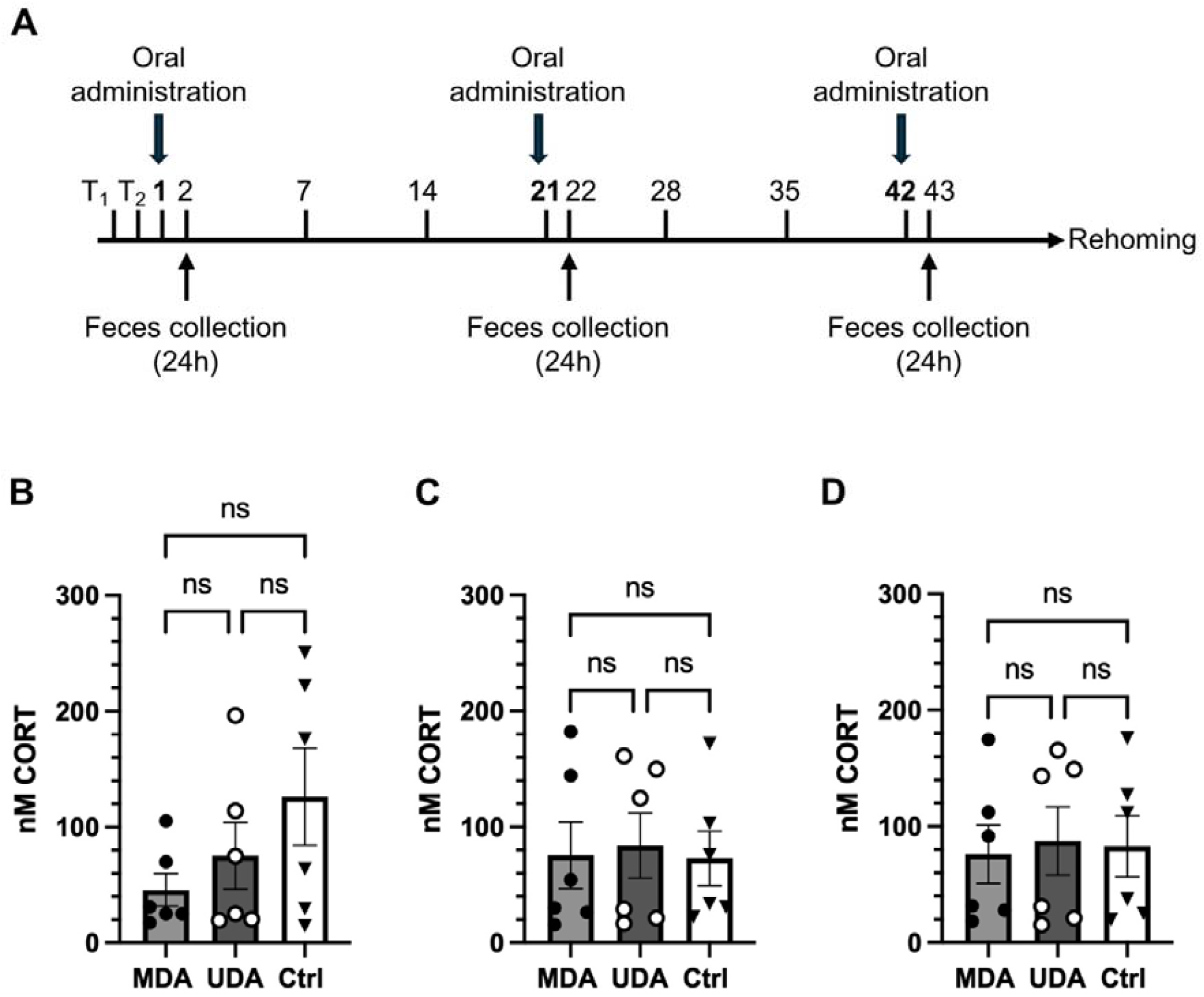
Levels of fecal corticosterone (CORT). A, design of the animal experiment showing the days of training (T1, T2), oral administration, feces collection and final rehoming of the animals. There were three groups, MDA, UDA and Ctrl (non-treated). Feces were collected over a 24-hour period after each oral administration procedure on days 2 (B), 22 (C), and 43 (D). Data represent the mean ± SEM of n=6; one-way ANOVA test, (ns) p> 0.05.

Moreover, we recorded the weights of the animals in each group for three consecutive days after each treatment, followed by weekly measurements to assess whether the UDA treatment influenced weight gain. Our results show a consistent weight gain across all groups and no significant differences between groups over the experimental period (Fig. 2).

**Figure 2.**
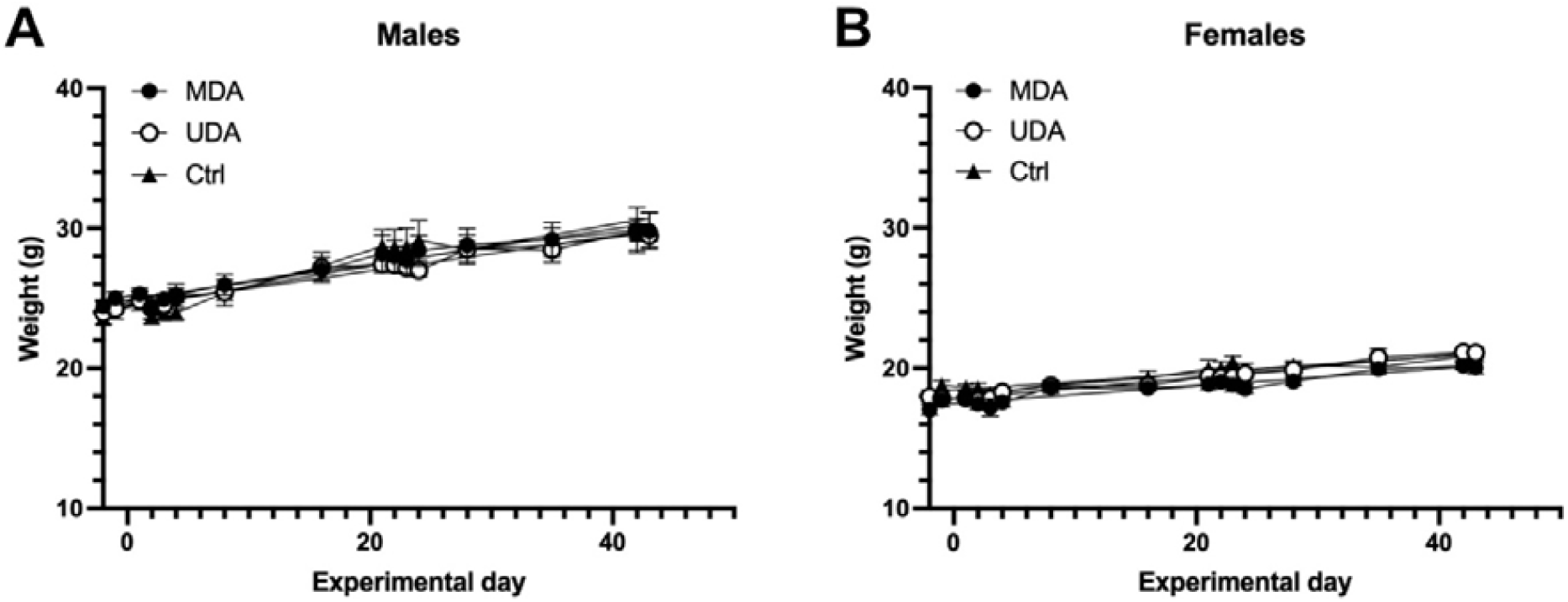
Oral treatment did not affect body weight gain. Body weights were measured during training sessions, before each treatment, on three consecutive days after treatment, and weekly between treatments. The data represent the mean ± SEM for n = 3 per sex and treatment. Linear regression analysis, p>0.05.

### Time of consumption using the UDA method

We recorded the time required to consume the agar units with UDA for each mouse and compared it with the time required by the MDA method. Since the mice were not isolated during the training sessions of the UDA method, the completion time was considered the same for all animals in the same cage. As depicted in Figure 3A, the consumption time on the first training day was significantly higher for UDA than for MDA, noting that MDA required full restraint of the mice. On the second training day, although still higher for UDA than for MDA, the consumption time significantly decreased from 20 minutes to about 1.5-2 minutes (Fig. 3B). For MDA, the average consumption time was roughly 0.5 minutes, but light restraint was still necessary. On the third day, which was considered the first administration day, the consumption time with the UDA method further decreased and was no longer significantly different from that with the MDA method, averaging about 1 minute (Fig. 3C and Video S1-S6). On the later administration days (days 21 and 42), we did not conduct new training sessions before administering UDA or MDA. We observed that the time for MDA did not change significantly, remaining between 30 and 60 seconds (Fig. 3D-E), although light restraint was still required in each case. While the mice still fully consumed the agar units, the time needed for UDA increased at day 21 (Fig. 3D), ranging from 3.6 to 27 minutes, and at day 42 (Fig. 3E), ranging from 3.75 to 31.6 minutes. Despite this longer completion time for UDA, fecal CORT levels remained low and were comparable to those in the MDA or non-administered control groups (Fig. 1C and D).

**Figure 3.**
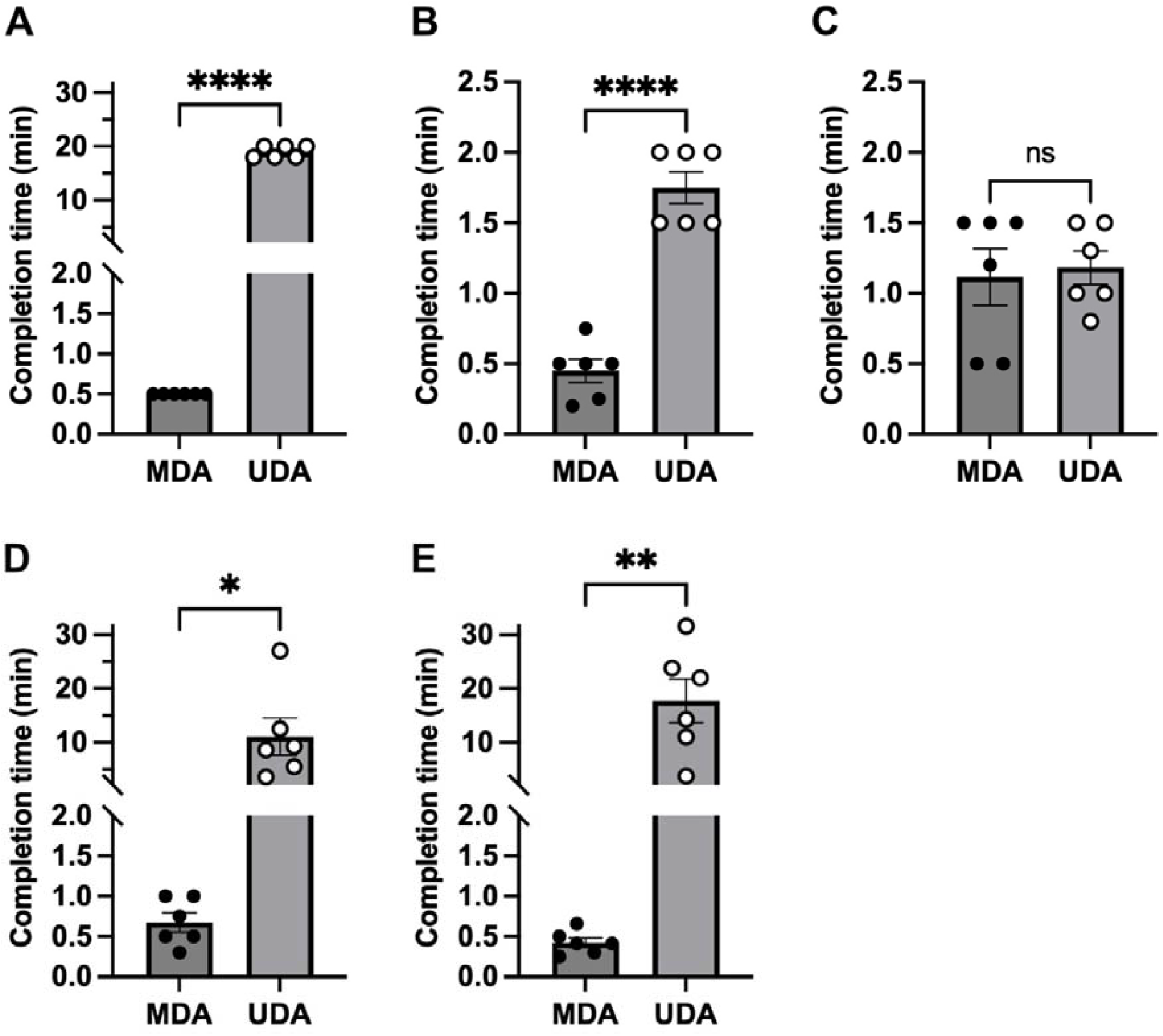
Time to completion for UDA vs MDA. The time required to complete each method was recorded in minutes. Plots for quantification of time (min) at (A) first training session; (B) second training session; (C) day 1 of oral administration; (D) day 21 of oral administration, and (E) day 42 of oral administration. The data represent the mean ± SEM of n = 6. Unpaired Student’s t-test, (ns) p>0.05, (*) p<0.05, (**) p<0.005, (****) p<0.0001.

## DISCUSSION

Oral administration is of extreme relevance in pharmacological studies, such as vaccine development, and oral gavage has traditionally been the standard method in animal experiments (17). Although oral gavage enables precise dosing, the disadvantages and complications associated with the technique, combined with growing demands for improved animal welfare, have prompted the exploration of alternative methods (4, 8, 9, 10, 11). With the aim of replacing oral gavage in mouse experiments and improving animal welfare, we developed a method for oral administration of substances in mice that, unlike other proposed methods, does not require fasting (9) or restraint (11, 14). To overcome the intrinsic neophobia of mice towards new food, with UDA, the animals require 2 days of training, down from 3 or 5 days reported in other methodologies (9, 14). The mice were trained in groups of three per cage without removing food or water. To adhere to the 3R (replace-reduce-refine) guidelines, we did not include an oral gavage group in these experiments, as it has already been demonstrated that oral administration of MDA results in significantly lower corticosterone levels than oral gavage (11). Our results revealed that the corticosterone levels were low and not significantly different from those obtained with the MDA technique (Fig. 1), indicating that UDA is indeed a low-stress method for oral administration in mice. For an experimental dose containing drugs, xenobiotics(18), or vaccines(19, 20, 21), the animals can be regrouped once they consume the agar unit. In our experiments, the time needed for consumption ranged from 0.8 to 1.5 minutes after training (Fig 3), increasing to a range from 3.6 to 31.6 minutes when dosing was done 21 or 42 days after the first dose. Despite these differences in time, the animals completely consumed the agar units, and it is worth noting that one advantage of this methodology is that multiple animals can be dosed simultaneously. The agar units in the present study had a volume of 400 μL, with the addition of condensed milk and bacon extract for palatability. This allows for more flexibility of the dosing volume, for example up from 35 to 55µl that can be administered by MDA to a group of mice of similar weight. In this study, we selected a dosing schedule that mirrors a typical vaccine trial; however, the approach is flexible and can be adapted for chronic administration, including daily dosing or multiple administrations per day. We conducted our experiments in C57BL/6 mice, and the timing of administration may vary if other strains are tested, as has been reported for MDA when comparing C57BL/6 and BALB/c (12).

## CONCLUSIONS

This study shows that UDA is a valid replacement for oral gavage in mice, thus contributing to animal welfare and experimental reproducibility. UDA is a low-stress, easy-to-adopt method for the oral administration of drugs, xenobiotics, or vaccines in mouse experiments. To overcome neophobia and to reduce the timing of administration, the method can be combined with two training sessions. On contrast to other methods described elsewhere, it requires no restraining or fasting and can be applied simultaneously to multiple animals. In addition, the agar formulation allows for doses containing non-water-soluble particles (e.g., xenobiotics, proteins, nanoparticles, viruses, spores or bacteria), it is palatable and has an attractive scent for the animals.

## LIST OF ABBREVIATIONS.

UDA: Unconstrained dosing agar.
MDA: Micropipette-guided drug administration.
FCM: Fecal corticosterone metabolites.
CORT: Corticosterone.

## DECLARATIONS

### Author’s contributions

ML, CE, CA conducted experiments and acquired data. CA, CE conceptualization, curation of data. CA, CE, CF wrote the manuscript. CA acquired funding.

### Data availability

Not applicable.

