## Supplemental Figure 1 for "Unconstrained dosing agar (UDA) Reduces Stress in Mouse Oral Administration"

**Supplementary Material**

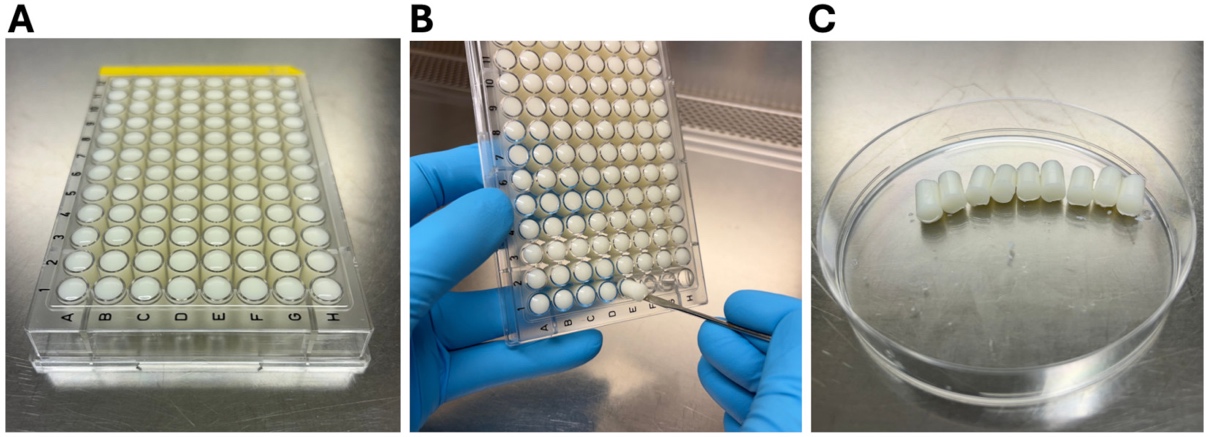

**Figure S1. Preparation of agar units for the UDA procedure**. (A) The formulation (see methods in the main text) was aliquoted into the wells of a sterile 96-well plate, then allowed to gel at room temperature. (B-C) The individual agar units were removed from the plate using a micro-spatula and stored at 4°C until further use, typically within 24h.

**Supplementary Videos**

**Video S1 - S6. UDA method on day 1 of administration**. Mice were isolated, and one agar unit was placed on the inverted mouse house. The video recording started when the cage was inserted into the rack (under 5 seconds after placing the agar unit), and the recording ended when the agar unit was fully consumed.
